# Quantifying virus load and characterising virus diversity in wildlife samples with target enrichment sequencing

**DOI:** 10.1101/2025.04.16.648874

**Authors:** Laura Bergner, Stefano Catalano, Jenna Nichols, Ana Da Silva Filipe, Xiaofei Cao, Daniel Mair, Andrina Nankasi, Moses Arinaitwe, Alfred Mubangizi, Oliver G Pybus, Claire Standley, Christina L Faust, Jayna Raghwani

**Affiliations:** School of Biodiversity, One Health and Veterinary Medicine, University of Glasgow, Glasgow, UK; MRC–University of Glasgow Centre for Virus Research, Glasgow, UK; Scotland Field Delivery, Animal and Plant Health Agency, Galashiels, UK; Genetic Design and Engineering Center, Department of Bioengineering, Rice University, Houston, USA; Vector Control Division, Ministry of Health, Kampala, Uganda; Pathobiology and Population Sciences, Royal Veterinary College, Hatfield, UK; Center for Global Health Science and Security, Department of Microbiology & Immunology, Georgetown University, Washington, DC, USA

## Abstract

Metagenomics is a powerful tool for characterising viruses, with broad applications across diverse disciplines, from understanding the ecology and evolutionary history of viruses to identifying causative agents of emerging outbreaks with unknown aetiology. Additionally, metagenomic data contains valuable information about the amount of virus present within samples. However, we have yet to leverage metagenomics to assess viral load, which is a key epidemiological parameter. To effectively use sequencing outputs to inform transmission, we need to understand the relationship between read depth and viral load across a diverse set of viruses. Here, using target enrichment sequencing, we investigated the detection and recovery of virus genomes by spiking known concentrations of DNA and RNA viruses into wild rodent faecal samples. In total, 15 experimental replicates were sequenced with target enrichment sequencing and compared to shotgun sequencing of the same background samples. Target enriched sequencing recovered all spike-in viruses at every concentration (10^2^, 10^3^, and 10^5^ ± 1 log genome copies) and showed a log-linear relationship between spike-in concentration and mean read depth. Background viruses (including *Kobuvirus* and *Cardiovirus*) were recovered consistently across all biological and technical replicates, but genome coverage was variable between virus genera and likely reflected the composition of target enrichment probe panel. Overall, our study highlights the strengths and weaknesses of using commercially available panels to quantify and characterise wildlife viromes, and underscores the importance of probe panel design for accurately interpreting coverage and read depth. To advance the use of metagenomics for understanding virus transmission, further research will be needed to elucidate how sequencing strategy (e.g. library depth, pooling), virome composition, and probe design influence viral read counts and genome coverage.

## INTRODUCTION

Studying virus transmission in wild animal populations is inherently challenging [1]. In contrast to humans and livestock, where longitudinal passive surveillance is routinely used to monitor disease dynamics, similar approaches are often impractical in wildlife populations. In particular, virus detection in wild animals is likely to be hindered by the absence of overt or recognisable clinical symptoms in natural hosts, reduced mobility and shelter-seeking behaviours in sick animals, and predation and scavenging of infected individuals [2]. These limitations underscore the need for improved tools and techniques to identify and track transmission of viruses in natural populations.

Virus metagenomic (‘metaviromic’) surveys have greatly accelerated the discovery of new viruses in all kingdoms of life, significantly expanding our knowledge of global virus diversity [3–7]. However, despite containing quantitative information on virus abundance within samples, metaviromic data has yet to be fully exploited for investigating the ecological [8] and epidemiological dynamics of viruses. Viral load is an important metric for understanding onward transmission [9, 10], identifying super-spreaders [11], and predicting clinical outcomes [12]. More recently, it has been shown that the distribution of virus quantitative PCR (qPCR) cycle threshold (Ct) values and viral reads within individuals can be used to understand population-level processes [13, 14]. Therefore, the ability to quantify abundance across a diversity of viruses would transform our ability to understand virus impacts on individuals and transmission within wildlife populations.

Traditionally, virus load is determined through antigen quantification, culture-based serial dilutions, or qPCRs that use specific primers [15]. Characterising virus loads across multiple viruses simultaneously has proved more difficult; optimizing multiplex PCR reactions requires time and resources and is not always possible for closely related viruses or under-characterised virus families [16]. Preliminary evidence suggests that metaviromics could help address this problem. For example, a comparison of qPCR Ct values with sequence read depth from individual-level host virome data from a mute swan population revealed a strong log-linear relationship, suggesting that read depth is a reliable proxy of virus load in wildlife [14]. Further support of the relationship between metrics of virus abundance in metaviromic data comes from controlled sequencing experiments, in which synthetic nucleotide sequences are added at known concentrations. These “spike-in” experiments show a positive correlation between virus read abundance and depth and virus load [17, 18].

One reason metaviromics remains underutilised for wildlife samples is that data generation is costly. Unbiased sequencing approaches often require high sequencing depth and can yield relatively small numbers of virus genomic data compared to background reads. One promising solution is target enrichment (also known as hybrid-capture) sequencing, which can simultaneously capture diverse virus genomes using custom-designed probe panels [19]. Target enrichment is well-suited for characterising low titre viruses and whole genome sequences of multiple virus species within samples with high proportions of background reads (i.e. faecal samples, host tissues) [19]. Furthermore, this method tolerates significant sequence divergence from target probes (10-20%; [20]) and has higher sensitivity, fewer sequencing errors, fewer false negatives (robust to mutations in primer sites), and fewer false positives (less likely to show low/uneven genome coverage) than amplicon based sequencing [21–23]. Custom probes have been used to characterise coronaviruses in bats [24–26], respiratory bacteria in chimpanzees [27], influenza viruses from waterfowl habitats [28], and to undertake more general virus surveillance from environmental sources [29]. All the work to date has used custom-developed panels, increasing the time and expertise needed to execute these protocols. Increasingly, commercial probe panels are becoming available that can target a broad range of microbes and could be deployed in small batches. Commercial probe panels would allow rapid assessment of low titre samples and require minimal troubleshooting, making them appealing for a diversity of research groups and sample types.

To understand the utility of target enrichment sequencing in quantifying and characterising known and unknown viruses in wild animal populations, we set-up a simple sequencing experiment where known concentrations of a mock viral community (comprising DNA and RNA viruses with diverse genome architectures) were spiked into a background matrix of RNA extracted from wild rodent samples. We also generated shotgun metagenomic data (i.e. without probe enrichment) from the same background samples. For spiked-in viruses (i.e. mock viral community), virus load was directly compared to read abundance and genome coverage. For viruses found in the background samples, the results were compared with data from shotgun sequencing and a custom qPCR. With this simple experimental design, we explored the benefits and challenges of using target enrichment sequencing to quantify and characterise viruses in wildlife populations.

## METHODS

### Sample collection and RNA extraction

As a part of a wider field study, we captured small mammals with live humane traps in July and August 2022 in small-holder agricultural fields in eastern Uganda (Mayuge and Buvuma Districts) and humanely euthanized target species via chloroform overdose (see Ethics Approval). We collected faeces from the rectum during post-mortem examinations and stored samples immediately in DNA/RNA Shield (DRS; Zymo Research Corp) at a 1:5 ratio by weight (e.g., 125mg faeces in 500μl DRS). Samples were stored initially at room temperature and then moved to -80°C after a maximum of 38 days at ambient temperature. For this study, we used three pooled faecal samples (M1, M2, and M3); each contained four individuals of *Mastomys erythroleucus* and pooled in equal volumes (Table S1). We included nuclease-free water controls in RNA extractions and for sequencing. We used ZymoBIOMICS RNA Miniprep Kit (Zymo Research Corp) to extract RNA from 150μl pooled faeces homogenate and eluted into 100 μl. We measured RNA concentrations using Qubit RNA HS Assay Kit on a Qubit 3 fluorometer (Thermo Fisher Scientific) and evaluated quality using a RNA 6000 Nano Kit on a 2100 Bioanalyzer (Agilient).

### Experimental design

We evaluated the performance of the Twist Comprehensive Viral Research Panel (CVRP; Twist Bioscience) in detecting and quantifying spike-in (known) and background (unknown) viruses in pooled rodent faeces. CVRP consists of 1,052,421 unique, 120 base pair (bp) probes that target 3,153 human and non-human associated viruses [30]. We selected a mock virome standard (Virome Nucleic Acid Mix; MSA-1008; ATCC:American Type Culture Collection) that includes six viruses with diverse genome compositions and structures (Human adenovirus 40, Human betaherpesvirus, Human respiratory syncytial virus, Influenza B virus, Mammalian othoreovirus 3,and Zika virus; Table S2). To evaluate recovery of viruses present at different initial abundances, we prepared experimental samples composed of three different background sample pools (M1, M2, and M3) and three dilutions of mock virome (10^2^, 10^3^, and 10^5^ ± 1 log genome copies). The spike-in concentrations chosen were based on a previous study evaluating sensitivity and specificity of the CVRP panel [17]. To evaluate protocol repeatability, we processed two technical replicates for each mock virome dilution in two background pools (M1 and M2), yielding a total of fifteen experimental samples (Table S3). Each background pool (M1-M3) was diluted to an initial concentration of 10ng/μl, to which an equal volume of the relevant dilution of virome mix was added. To monitor contamination during library preparation, a nuclease-free water negative control was processed alongside experimental samples.

### Library preparation for target enrichment sequencing

We prepared libraries using workflows for the Total Nucleic Acids Library Preparation EF v2.0 and Target Enrichment Standard Hybridization v1.0 Kits (Twist Bioscience). We followed all manufacturer’s instructions, except for modifications to first strand cDNA synthesis for some samples (more details in Supplementary Material). We diluted experimental samples to a concentration of 3.3 ng/μl of background RNA and volume of 15μl. We performed first strand cDNA synthesis using ProtoScript II (New England Biolabs) for samples with background M1 and M2, while SuperScript II (Thermo Fisher Scientific) was used for samples with background M3 and the negative control. We sequenced pools on an Illumina NextSeq500 using a Mid Output 300-cycle cartridge, 2 x150bp with a target of 16M reads per sample at the MRC-University of Glasgow Centre for Virus Research.

### Library preparation for shotgun sequencing

We used Glasgow Polyomics’ sequencing service for shotgun sequencing. Samples were prepared using the QIAseq FastSelect Epidemiology Kits (Qiagen) to remove rRNA and then the NEBNext® Ultra™ II Directional RNA Library Prep Kit for Illumina (New England Biolabs) to create libraries. They sequenced samples on an Illumina NextSeq2000 using a P3 200-cycle cartridge, 2 x100bp with a target of 5M reads per sample.

### Quantification of background *Kobuvirus* using qPCR

To assess the capture efficiency of the CVRP for quantifying viruses present in the rodent sample background (i.e. viruses that weren’t spike-in), we developed a qPCR assay for a rodent *Kobuvirus* detected in experimental samples (see Results). Primers and probes for the qPCR assay were designed for the *Kobuvirus* polyprotein gene using the Integrated DNA Technologies (IDT)PrimerQuest tool and purchased from IDT (Table S4). We generated cDNA from 300 ng extracted RNA from M1 and M2 with the SuperScript III First-Strand Synthesis System (Invitrogen) and random hexamers, which was then stored at -20°C.

We performed qPCR reactions on cDNA using IQ Supermix (BioRAD) and the newly designed *Kobuvirus* primers/probe on an AriaMx Real time PCR Instrument (Agilent Technologies) with the following cycling conditions: 5 minutes at 95°C, and 59 cycles of 5 seconds at 94°C and 45 seconds at 60°C. We used serial 10-fold dilutions of extracted plasmid DNA containing a positive control fragment to determine amplification efficiency (see Supplementary Information). We did not use template controls as negative controls, and we ran both experimental samples and positive controls in triplicate. We generated a standard curve with positive control dilutions from 10^-2^ - 10^-8^, which demonstrated that the qPCR had an acceptable R^2^ value of 0.997 and efficiency of 108%.

### Bioinformatics

Raw read data were processed using a modified metagenomic workflow[14, 31], which included adapter removal, filtering reads with quality scores ≥30 and read length ≥45bp using CUTADAPT 1.18 [32]. Cleaned paired-end reads were de novo assembled into contigs using MEGAHIT 1.2.8 [33]. Assembled contigs were classified taxonomically using DIAMOND 0.9.22 [34] and the NCBI non-redundant database (downloaded December 2023). Viral contigs were identified based on e-value scores < 10^-5^. To determine read depth and coverage of spike-in viruses, reads were mapped against reference genomes provided by ATCC using bowtie2 under the “very-sensitive” mode [35]. Coverage and depth were calculated before and after removing PCR duplicate reads using samtools [36]. Code is available on the public github repository: https://github.com/cfaustus/te_ug_rodents and raw read data from this study is available in the Sequence Read Archive (SRA) under BioProject accession number PRJNA1250535.

For background viruses, assembled viral contigs from shotgun sequence data were used as reference genomes to obtain estimates of read depth and coverage for both the target enrichment and shotgun sequence datasets. The same bioinformatic pipeline was used to re-analyse published data from a study which spiked-in the same mock virome into a synthetic human sample background, and also used CVRP for target enrichment [18].

## RESULTS

We examined the efficacy of a commercially available target enrichment probe panel in characterising unknown viruses and in quantifying abundances of spike-in viruses at known concentrations in wild rodent faecal samples. On average, we obtained 10,667,076 (range 565,371 - 53,224,935) cleaned reads per sample, while the negative control contained 21,999 reads (none of which were viral contigs). Samples were processed in two hybridization pools. Pools were similar in sequencing depth (Pool(P)1 mean depth: 10,411,807; P2 mean depth: 9,591,710) (Figure S1). Three experimental samples with the M3 sample background were excluded from further analyses as these were processed differently during cDNA synthesis and lacked technical replicates (see Methods). Although not directly comparable due to the presence of spike-in viruses in target enrichment samples, shotgun sequencing of the two sample backgrounds yielded lower proportions of viral reads (M1: 1.2%, M2: 0.5%) compared to target enriched samples with 10^2^ spike-in viruses (M1: 2.7%, M2: 5.6%) (Table S3).

### Target enrichment sequencing of spike-in viruses

We detected all six of the spike-in viruses in replicated sample backgrounds M1 and M2, and at each viral load (10^2^, 10^3^, and 10^5^) (Figure 1). For each spike-in virus, we observed an increase in read count and read depth with increasing viral loads, which was consistent regardless of method used for normalisation and whether reads were deduplicated or not (Figure S2). Log-linear models showed a similar effect of viral load on mean read depth across viruses, with only minimal variation in the regression slope (range 0.91 - 0.95) and R^2^ (range 0.88 - 0.92) across viral taxa (Figure 1). After normalising for genome length, we observed relatively lower read depths for DNA viruses (Adenovirus and Betaherpesvirus), and similar read depths for RNA viruses, with the exception of Mammalian Orthoreovirus, which had higher read depth (Figure S2). There was some evidence of variability between hybridization pools in terms of viral read count and mean read depth (Figure S1). Trends in viral read count and the proportion of viral reads were relatively consistent between sample backgrounds (Figure S3).

**Figure 1.**
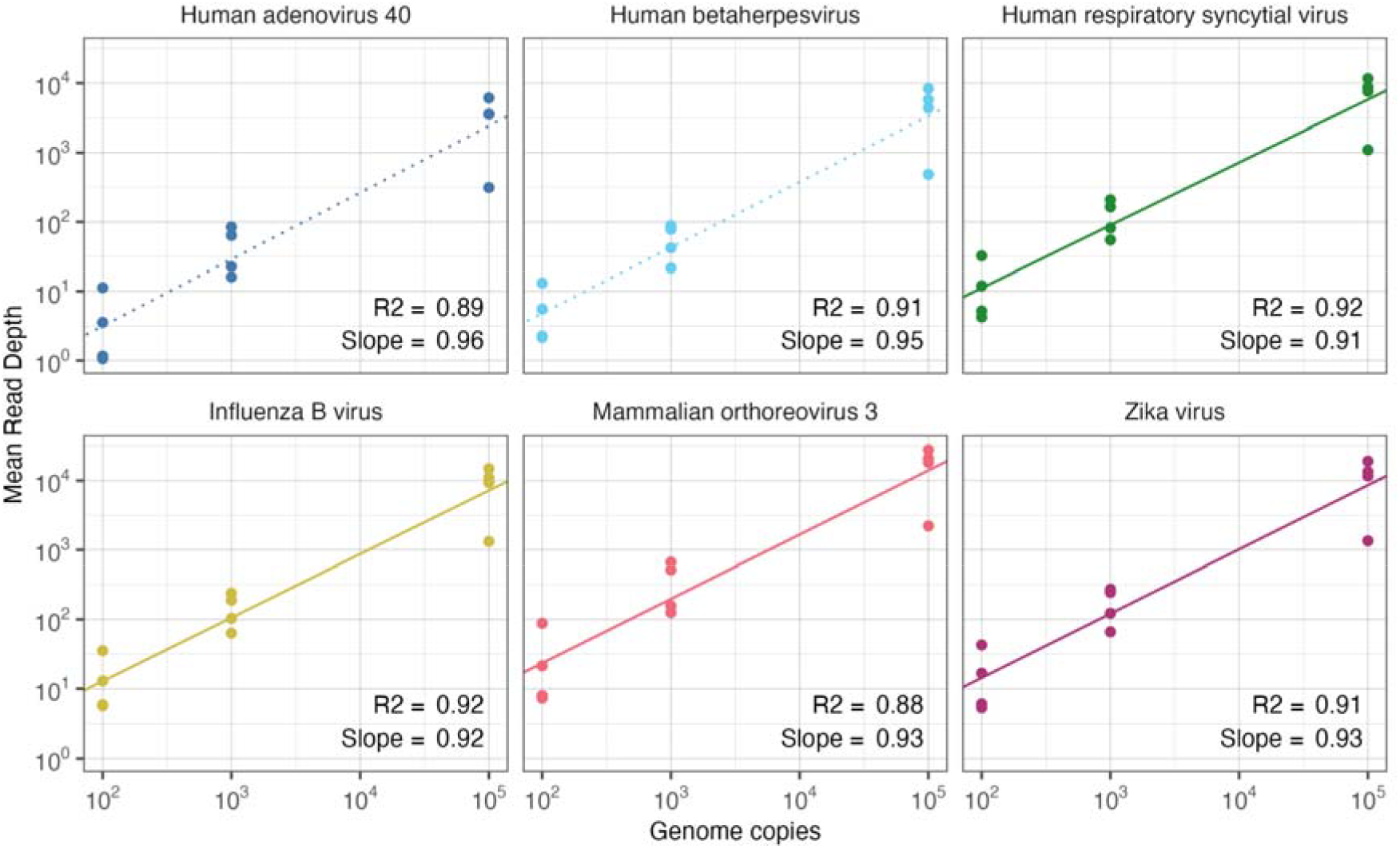
Relationship between spike-in viral load and mean read depth per virus. Deduplicated read depths are shown for M1 and M2 backgrounds. Linear models are fit to virus species. Dotted lines are used for DNA viruses and solid lines are used for RNA viruses.

Genome coverage increased with spike-in viral load (Figure 2). We recovered full genome coverage for all six viruses at higher viral loads (10^3^, 10^5^) in all replicates. At lower concentrations, genome coverage was affected by virus species and sample background, with relatively lower coverage observed for DNA viruses and for the M1 sample background (Figure 2, Figures S4-S9). Compared to the other spike-in viruses, Human Adenovirus exhibited lower read depth and more sparse genome coverage, particularly at lower viral loads (Figures 2, S2, and S4). To understand this observation, we reanalysed data from another study that also evaluated CVRP with the same mock virome standard [18], which similarly found lower depth and coverage for Human Adenovirus (Figures S10-S12).

**Figure 2.**
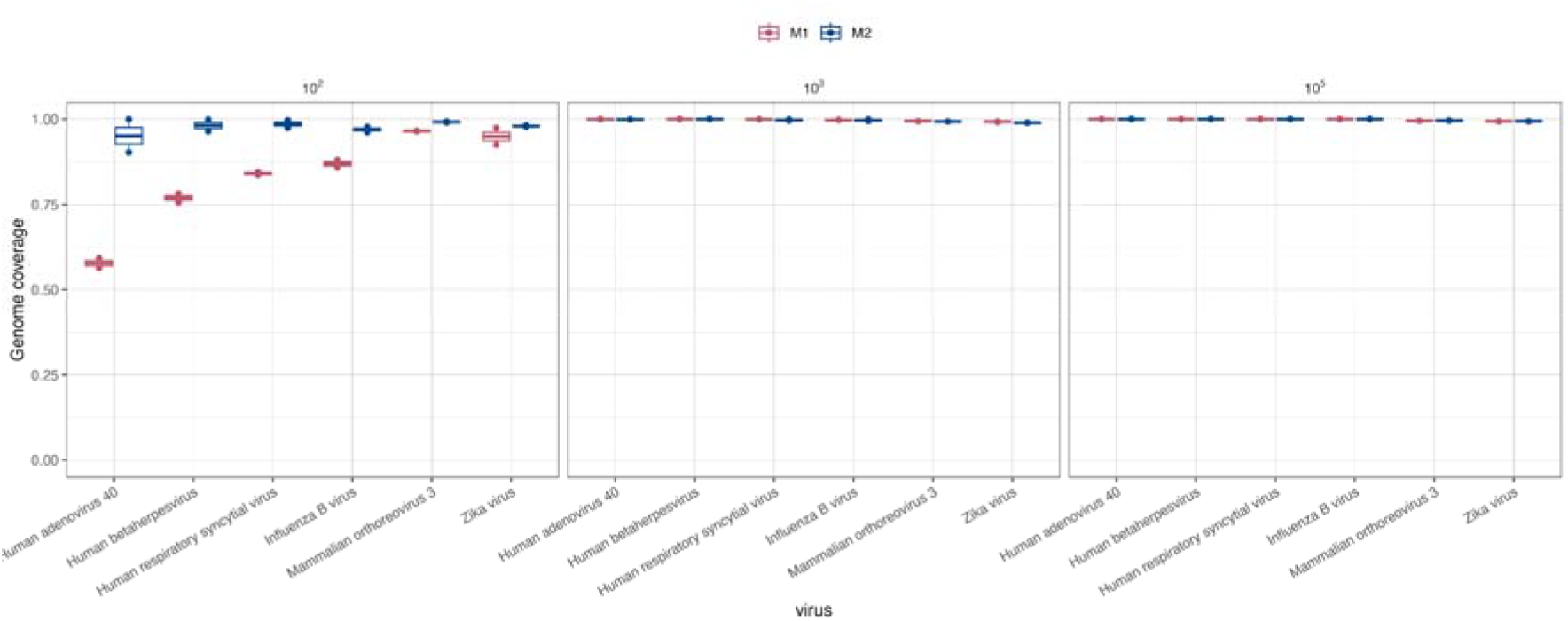
Genome coverage of spike-in viruses across spike-in viral loads. Boxplots are colored by background sample (M1 or M2) and plots are arranged left to right with increasing spike-in loads.

### Target enrichment sequencing of background viruses

We detected several non spike-in viruses in our samples, including two picornaviruses (genera *Cardiovirus* and *Kobuvirus*) and one circovirus (genus *Cyclovirus*). *Cardiovirus* and *Kobuvirus* were found in both sample backgrounds and in shotgun sequence data, and are hereafter referred to as ‘background viruses’. We observed lower read counts (Figure 3), read depths (Figure S13), and genome coverage for *Cardiovirus* than for *Kobuvirus* (Figure S14). We found that reads were clustered in certain regions of the Cardiovirus genome for target enrichment sequencing, but not shotgun sequencing (Figure S15).

**Figure 3.**
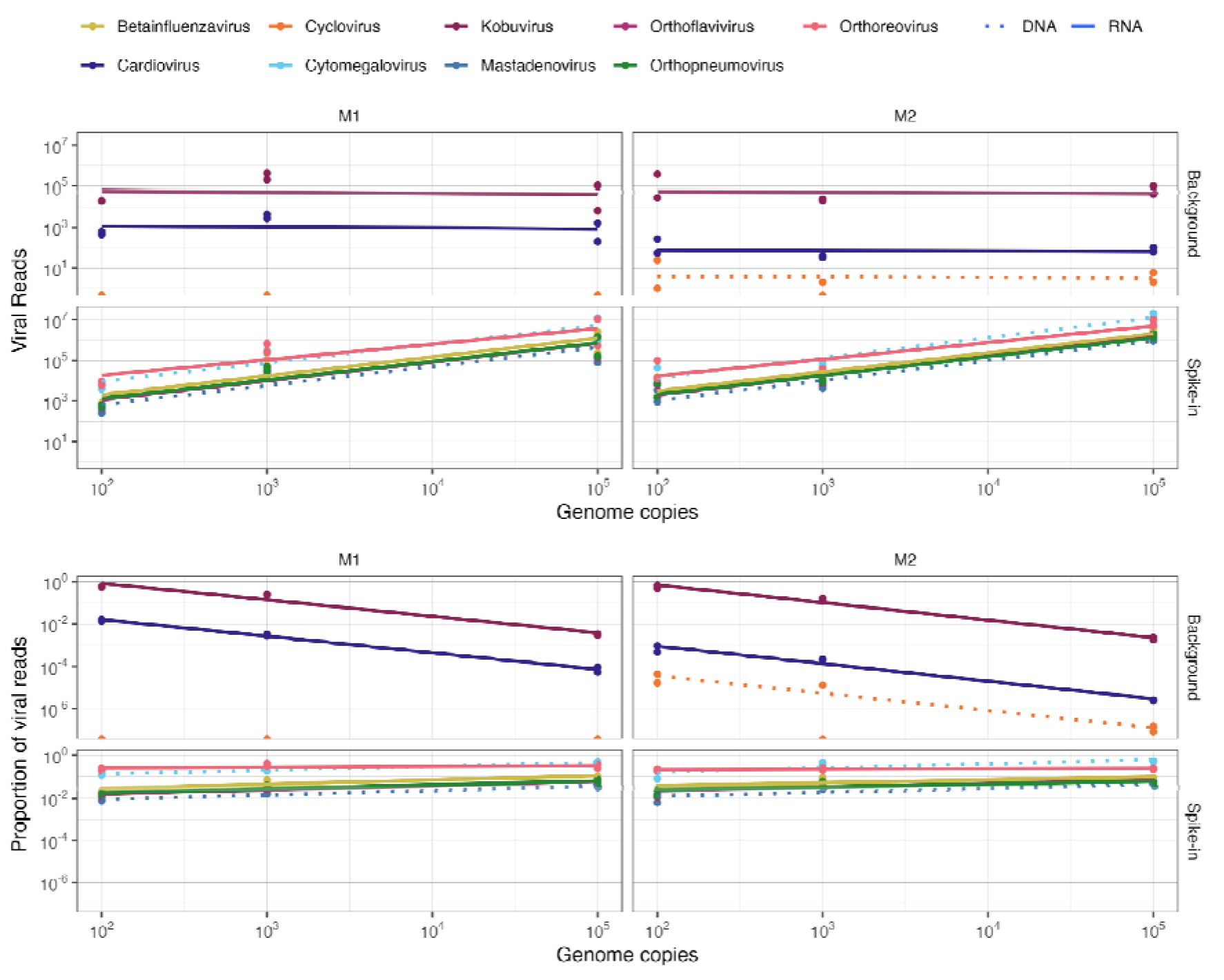
Recovery and read depth of spike-in and background viruses across technical and biological replicates. Lines are predictions from linear models fit to genera. The linear fit to background viruses is poorer (i.e. more variation) compared to spike-in viruses for viral reads.

The read count of background viruses did not notably vary with increasing spike-in viral loads (Figure 3). However, the proportion of both total and viral reads, which include background viral reads, decreased as spike-in viral loads increased (Figure 3). We further examined the impact of spike-in viral loads on background viruses by focusing on the rodent *Kobuvirus*, which was detected in all experimental samples (Figure 3). Spike-in viral loads did not affect read depth and genome coverage, which were consistently high across all replicates (Figures S13, S14). Instead, sample background appeared to be more important in determining the overall genome coverage and coverage per site. Specifically, samples with M2 background consistently exhibited higher coverage for *Kobuvirus* in both target enrichment and shotgun sequence datasets (Figure S14, S16). A similar but opposing pattern was observed for *Cardiovirus*, with coverage and depth generally higher in M1 background (Figures S13-S15).

Rodent *Kobuvirus* was also detected in shotgun metagenomic data from the same background samples. We used qPCR Ct values and read depth from shotgun metagenomic data to determine if target enriched samples had comparable viral loads. Mean read depth was higher for M2 background than M1 in the shotgun metagenomic data (330.6 compared to 145 for deduplicated data), but there was no clear relationship between mean read depth and sample background in the target enrichment dataset (Figure 3, S16). Compared to shotgun data, target enrichment samples had higher per site coverage across samples in some *Kobuvirus* genomic regions (Figure S16). qPCR data showed that the pools differed significantly in mean virus load (Welch t-test t=6.29; p=0.01), with lower Ct values indicating higher viral load for M2 (mean Ct 26.7+/-0.12) compared to M1 background (mean Ct 28.3+/-0.4).

## DISCUSSION

Our study demonstrates that target enrichment sequencing can quantify and characterise spike-in and background viruses in wild rodent samples. Spike-in viruses were recovered at all tested concentrations, and we found a strong positive correlation between virus loads and mean read depth. Background rodent viruses were also enriched, but genome coverage was generally lower compared to spike-in viruses. Overall, our findings suggest that commercial target enrichment probe panels can be used to process wildlife samples, but custom designed probes are likely required for more precise quantification of virus load and higher genome coverage.

Although spike-in viruses were successfully recovered, mean read depth and genome coverage for certain viruses notably varied, particularly at lower viral loads. The variation in read depth between dsRNA, ssRNA, and DNA viruses suggests that genome structure influences the efficacy of target enrichment, which has been previously observed by other studies [17, 18]. This pattern could be related to cDNA synthesis and probes for both strands of dsRNA viruses. Differences in tiling coverage across viral genomes could also contribute to variation in capture efficiency between viral strains. The discernible lower depth and coverage of Human adenovirus at low spike-in load is consistent with a previous study that evaluated Twist CVRP with ATCC virome mix [18], suggesting this observation is unlikely caused by any laboratory or bioinformatic pipeline. Instead, it indicates some unknown interaction between the probe panel and spike-in adenovirus genome sequence. Altogether, our results highlight the importance of understanding the probe panel design, e.g. total number of probes, tiling density, relative concentration of probes across the virus genome, when evaluating virus loads with genomic data.

We recovered both DNA (*Cyclovirus*, ssDNA) and RNA (*Kobuvirus* and *Cardiovirus*, ssRNA) viruses from the background samples in all replicates. Given that CVRP primarily targets human viruses, the detection and coverage of background viruses will likely depend on the degree of sequence similarity to probes and how well the genetic diversity is represented by the probe panel. For example, while both *Kobuvirus* and *Cardiovirus* belong to the family Picornaviridae, differences in their detectability suggests that *Kobuvirus* genetic diversity is better accounted for by the probe panel compared to *Cardiovirus*. This is despite both genera having a similar number of virus representatives within the CVRP: *Kobuvirus* representatives (n = 31) compared to *Cardiovirus* representatives (n = 26). The higher genome coverage of *Cardiovirus* observed in the shotgun data supports this idea, and strongly suggests that the lower coverage of *Cardiovirus* in target enrichment data is likely due to probe mismatch. Both shot-gun sequencing and qPCR demonstrated higher levels of Kobuvirus in M2 compared with M1, but similar mean read depths were observed among target enriched sequencing replicates. This discrepancy may be due to the decision to equimolar all background samples prior to target enrichment library preparation. Beyond the technical aspects of probe panel design, our findings underscore that we also need to carefully consider the workflow and composition of the probe panel when using this sequencing approach for virus discovery in wildlife samples.

Our study demonstrates that multiple viruses can be quantified simultaneously with target enrichment sequencing, which represents a huge potential in understanding wildlife virus communities. Virus genomes with matches in the probe panel (i.e. Influenza B) and divergent from the probe panel (i.e. *Kobuvirus*) were sequenced at read depths comparable to spike-in concentrations and evaluation of genome copy number through custom qPCR. In theory, at higher concentrations of viruses, there will be competition for probe hybridization and sequencing depth, which would complicate multi-species evaluation. However, we did not see any plateauing of mapped read depth at higher concentrations; this may be due to the limited serial dilutions (three) or that concentrations were not yet at saturation levels. The differences observed between DNA and RNA viruses suggest there may be predictable effects of genome structure on read recovery and could be taken into account. Additional studies will be needed to determine if interpreting viral reads should statistically account for observed variability in the future (e.g. number of probes per viral genome, genome composition, reads per pool). In the future, artificial spike-ins could be used to help normalize between individual samples and improve comparability of read depths. There is also scope for future work to evaluate how analytical choices, including methods for normalization, affect the outcomes and interpretability of these data.

## CONCLUSION

Our study further supports the use of viral reads from target enrichment sequencing to assess virus load. The ability to quantify viruses is essential for understanding processes that shape the temporal and spatial patterns of virus shedding in wildlife populations. These methods can be scaled to larger studies to map virus load across populations and ascertain when and where transmission risk is elevated. This approach is valuable for understanding virus circulation in natural populations and/or assessing zoonotic spillover risk.

## Acknowledgements

We appreciate the field team members from the Vector Control Division, Ministry of Health Uganda, that collected samples. We also appreciate the community leaders, village health teams, and District Vector Control Officers (Henry Magala and Juma Nabonge) for facilitating access to sites. We also acknowledge the Centre for Virus Genomics team and Glasgow Polyomics for advice and assistance with sequencing.

## Funding statement

The study was supported by a US-UK EEID Biotechnology and Biological Sciences Research Council and Medical Research Council grant (BB/Y006879/1). Fieldwork was supported by the Natural Environment Research Council (NE/V014730/1). Laboratory work was supported UNESCO-L’Oreal For Women in Science and the Defense Threat Reduction Agency, United States Department of the Defense (HDTRA12110028) (the content of the information does not necessarily reflect the position or the policy of the federal government, and no official endorsement should be inferred). Shotgun sequencing was supported by Wellcome Trust Institutional Strategic Support Fund (204820/Z/16/Z).

## Conflict of interest statement

Authors declare there is no conflict of interest.

## Data summary

The authors confirm that supporting data, code and protocols are publicly available or available within the article, supporting information and associated links. Code is available on github https://github.com/cfaustus/te_ug_rodents. All raw sequencing data have been deposited in SRA under BioProject accession number PRJNA1250535.

## Impact statement

There is an increased interest in characterizing and quantifying viruses in wildlife populations. We performed an experiment to determine if virus load could be quantified using target enrichment sequencing. We demonstrated that known quantities of viruses with diverse compositions and structures could be recovered and quantified with target enrichment sequencing. Additionally we show that background viruses that are not directly in the probe panel can be recovered and quantified. This study provides validation for using target enrichment for not only describing virus species, but also quantifying the virus load.

## Ethics approval

Small mammal trapping and sampling protocols were reviewed and approved by the Vector Control Division Research Ethics Committee (VCDREC/095), Ugandan National Council for Science and Technology (NS/622) and the University of Glasgow’s School of Veterinary Medicine Ethics (27a/18).

## Author contributions

LB: Data curation, Formal Analysis, Writing-original draft preparation; SC: Methodology, Investigation, Writing – review & editing; JN: Methodology, Writing – review & editing; ADSF: Conceptualization, Investigation, Writing – review & editing; XC: Investigation AN: Investigation; MA: Investigation; AM: Supervision; OP: Supervision, Funding Acquisition, Writing – review & editing; CS: Supervision, Funding Acquisition, Writing – review & editing; CF: Conceptualization, Investigation, Supervision, Funding Acquisition, Writing-original draft preparation; review & editing; JR: Conceptualization, Investigation, Software, Supervision, Funding Acquisition, Visualization, Writing-original draft preparation; review & editing

## Notes

https://www.ncbi.nlm.nih.gov/sra/PRJNA1250535

### Competing Interest Statement

The authors have declared no competing interest.

