## Supplementary Information for "Quantifying virus load and characterising virus diversity in wildlife samples with target enrichment sequencing"

Figure S6. Genome coverage of human respiratory syncytial virus ………….....….9

Figure S7. Genome coverage of Influenza B virus…………………………………...10

Figure S8. Genome coverage of Mammalian orthoreovirus 3……………………...11

### SUPPLEMENTARY METHODS

#### RNA Extraction

RNA extractions were performed using ZymoBIOMICS™ RNA Miniprep Kit (Zymo Research Corp). We used pooled faeces in a buffer comprising equal volumes of four individual samples that were initially stabilized in DNA/RNA Shield (Zymo Research Corp) in the field. 150µl of pooled faecal samples were added into a bead-bashing tube containing 150µl molecular grade water and 150µl DNA/RNA Shield 2X concentrate. These were immediately bead-beaten using Precellys Evolution Touch (Bertin Technologies SAS) for two cycles of 15 sec each at 5,000 rpm and 10 sec rest between cycles. Homogenates were centrifuged at 16,000g for 1 min, after which 250µl supernatant was transferred into a new 2 ml tube - the remainder of the extraction followed the manufacturer's protocol, including DNase treatment.

**Table S1**. Individuals used in pooled RNA samples

All samples are from Uganda. Samples from Buvuma District are from the main Buvuma Island on Lake Victoria. Mayuge District is east and adjacent to Buvuma District. The two Mayuge pools are from populations >30 km apart.

| **Background sample** | **Unique rodent ID** | **Location** | **Species** | **Sex** |
|---|---|---|---|---|
| M1 | RBB387 | Buvuma District | *Mastomys erythroleucus* | M |
| M1 | RBB404 | Buvuma District | *Mastomys erythroleucus* | F |
| M1 | RBB405 | Buvuma District | *Mastomys erythroleucus* | F |
| M1 | RBK381 | Buvuma District | *Mastomys* species | F |
| M2 | RWA311 | Mayuge District (West) | Mastomys species | M |
| M2 | RWA320 | Mayuge District (West) | Mastomys species | M |
| M2 | RWA339 | Mayuge District (West) | Mastomys species | M |
| M2 | RWA342 | Mayuge District (West) | Mastomys species | F |
| M3 | RKB456 | Mayuge District (East) | *Mastomys erythroleucus* | M |
| M3 | RKB459 | Mayuge District (East) | Mastomys species | F |
| M3 | RNA478 | Mayuge District (East) | *Mastomys erythroleucus* | M |
| M3 | RNA479 | Mayuge District (East) | *Mastomys erythroleucus* | M |

#### Details of experimental design and preparation of target enrichment libraries

<u>cDNA synthesis:</u>

We performed first strand cDNA synthesis using two separate protocols. The ProtoScript II protocol, Twist's recommended protocol, was applied to all M1 and M2 samples. Briefly, 5µl random primer mix (New England Biolabs) was combined with 50 ng of background pool (diluted to 15ul). The mixture was incubated at 95°C for 5 min then placed on ice. 25µl ProtoScript II reaction mix and 5µl ProtoScript II enzyme mix (New England Biolabs) were added to samples for a total volume of 50µl. The reaction was incubated at 25°C for 5 min, 42°C for 60 min, 80°C for 5 min, followed by 4°C hold. The SuperScript II protocol was applied to all M3 samples and the negative control. Briefly, 1.5 µl dNTP mix at 10mM (New England Biolabs) was combined with 1.5µl random hexamers at 50µM (Invitrogen), 50ng background pool (diluted to 15µl) or water negative control from extractions, and 2 µl nuclease free water. The reaction was incubated at 65°C for 5 min then placed on ice. Following this, 10µl 5X SuperScript II first-strand buffer (Invitrogen), 5 µl 100mM DTT, 2.5µl RNaseOUT (Invitrogen), 2.5µl SuperScript II Reverse Transcriptase, and 10µl nuclease free water were added to samples. Samples were incubated at 25°C for 10 min, 42°C for 50 min, 70°C for 15 min, followed by 4°C hold.

We performed second strand cDNA synthesis using the NEBNext Ultra II Non-Directional Second Strand Synthesis Module (NEB) for all samples. After cDNA synthesis, we measured sample concentrations using a Qubit dsDNA High Sensitivity Assay Kit (Invitrogen) and evaluated size distribution and fragmentation for a subset of samples using a TapeStation High Sensitivity D1000 Assay (Agilent). Double stranded cDNA samples were stored overnight at 4°C.

**Table S2**. Virus in mock virus community

Virus species present in the Virome Nucleic Acid Mix (https://www.atcc.org/products/msa-1008) and their characteristics. Viruses represent a diversity of structures and compositions and genomes range in size from 11-230 kbp.

| **Virus** | **Composition** | **Structure** | **Genome size (kbp)** |
|---|---|---|---|
| Human adenovirus 40 | dsDNA | nonenveloped | 36 |
| Human herpesvirus 5 | dsDNA | enveloped | 230 |
| Human respiratory syncytial virus | ssRNA | enveloped | 15 |
| Influenza B virus B/Florida/4/2006 | ssRNA | enveloped | 14 |
| Mammalian orthoreovirus 3 | dsRNA | nonenveloped | 23 |
| Zika virus | ssRNA | enveloped | 11 |

Target Enrichment:

We performed fragmentation, end-repair and A-tailing according to the manufacturer's instructions, with fragmentation time set at 30°C for 15 minutes (desired insert size 275-350 bp). We ligated Twist Universal Adapters and cleaned products using DNA Purification Beads. Finally, we amplified libraries using Twist UDI primers with twelve PCR cycles, followed by cleanup with DNA Purification Beads. We quantified final pre-capture libraries using a Qubit dsDNA Broad Range Assay Kit and evaluated size distributions using a TapeStation High Sensitivity D1000 Assay. We stored pre-capture libraries at -20°C.

We performed target enrichment (Twist Bioscience) according to the manufacturer's instructions. Briefly, we combined 200 ng of each of the amplified, indexed libraries prepared into two 8-plex pools (Table S3). We concentrated pooled libraries following the Alternate Pre-Hybridization DNA Concentration Protocol and hybridised capture probes to pools, incubating at 70°C for 16 hours. We then amplified post-capture pools with 8 PCR cycles and cleaned with DNA Purification Beads.

**Table S3**. Individual samples for target enrichment sequencing

Individual sample characteristics - technical and biological replicates are randomly assigned to the target enrichment pools (P1, P2). All reads reported are deduplicated.

| Sample ID | Background sample | Spike-in viral load | Target Enrichment Pool | Number of paired-end reads (QT) | % viral reads |
|---|---|---|---|---|---|
| A | M3 | 10^3 | P2 | 1087372 | 2.870 |
| B | M1 | 10^2 | P1 | 717265 | 2.591 |
| C | M2 | 10^5 | P1 | 24205188 | 50.199 |
| D | M2 | 10^3 | P1 | 775693 | 9.845 |
| E | Neg_control | 0 | P2 | 21999 | 0.845 |
| F | M3 | 10^5 | P2 | 12428910 | 16.115 |
| G | M1 | 10^2 | P1 | 565371 | 2.881 |
| H | M2 | 10^2 | P2 | 3708723 | 7.418 |
| I | M3 | 10^2 | P2 | 3977262 | 53.314 |
| J | M2 | 10^3 | P1 | 694613 | 12.918 |
| K | M2 | 10^5 | P1 | 53224935 | 38.398 |
| L | M1 | 10^5 | P1 | 2334898 | 47.816 |
| M | M1 | 10^3 | P2 | 2678683 | 15.375 |
| N | M1 | 10^5 | P2 | 41327620 | 35.894 |
| O | M2 | 10^2 | P1 | 776496 | 3.837 |
| P | M1 | 10^3 | P2 | 11503116 | 7.013 |

<u>Generation and amplification of Kobuvirus qPCR positive control:</u>

We generated a qPCR positive control by first synthesising cDNA with a gene specific primer (Table S4) and the SuperScript III First-Strand Synthesis System (Invitrogen). The positive control was generated using RNA from an experimental pool (ME_P4) which was not used for metagenomic sequencing or target enrichment sequencing. PCR reactions to amplify the positive control were carried out with Taq DNA Polymerase (Invitrogen) and primers synthesised by Integrated DNA Technologies (Table S4) using the following conditions: 10 min at 94°C for 1 cycle, 30 sec at 94°C, 45 sec at 55 °C, and 1 min at 72°C for 40 cycles, 10 min at 72°C for 1 cycle, hold at 12°C. The PCR product was visualized by 2% Agarose gel with SYBR Safe DNA Gel Stain (Invitrogen) and cloned into pCR™4-TOPO TA vector as outlined by manufacturer's instructions, using the TOPO TA Cloning® Kits for Sequencing (Invitrogen). Whole colony PCR was performed on the selected transformants to specifically amplify the insert region using universal primers M13 forward (-20) primers and M13 reverse primer. Plasmid DNAs were extracted from overnight cultured positive colonies using Monarch Spin Plasmid Miniprep Kit (New England Biosciences). The insertion of positive control fragments was confirmed using Sanger sequencing (Genewiz).

**Table S4.** Description of *Kobuvirus* qPCR primers and probes.

| Primer/Probe name | Description | Sequence |
|---|---|---|
| KobuV_HT_S1F_2058 | qPCR Forward Primer | CCCTTCTCCTTTGTGCGTGCTTAC |
| KobuV_S5R_2240 | qPCR Reverse Primer | GGAACAGGAGAGGGAGGT |
| KobuV_Probe1_2107 | qPCR Probe | 5’ 6-FAM/ ACAACACCCACTCCATGTGGAACTGCGGGT /3’ BHQ-1 |
| KobuV_PCRT_S4R_934 | Gene specific reverse transcription primer for positive control | ATGGACTTGGGCAGGAATG |
| KobuV_PCEP_S1F_308 | PCR forward primer for Positive control | CCAGATTTCCCTATTTCCACAGA |
| KobuV_PCEP_S1R_699 | PCR reverse primer for Positive control | ACTTGGAGGTAGAAGGTCCA |

### SUPPLEMENTARY FIGURES

#### Figure S1. Library capture pool impacts

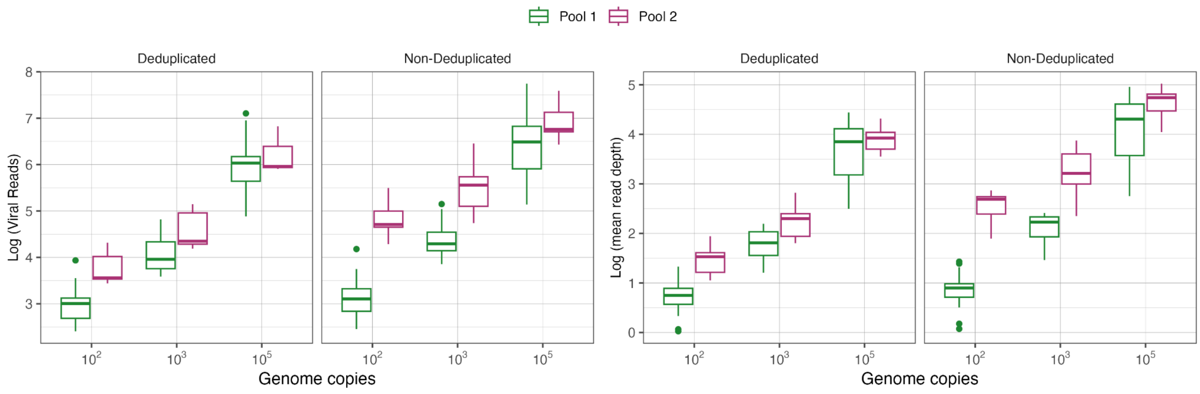

#### Figure S2. Spike-in virus quantities with different normalization methods

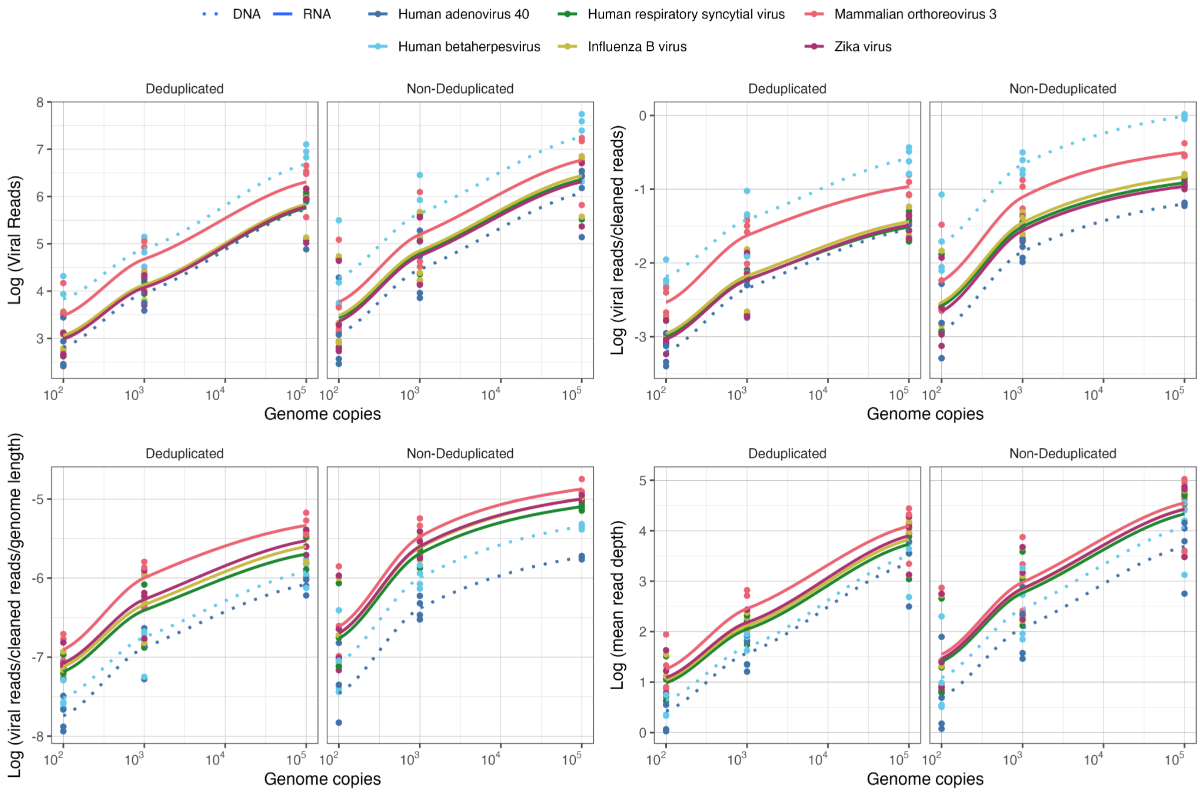

**Figure S3. Total reads, total viral reads and proportion viral reads by background**

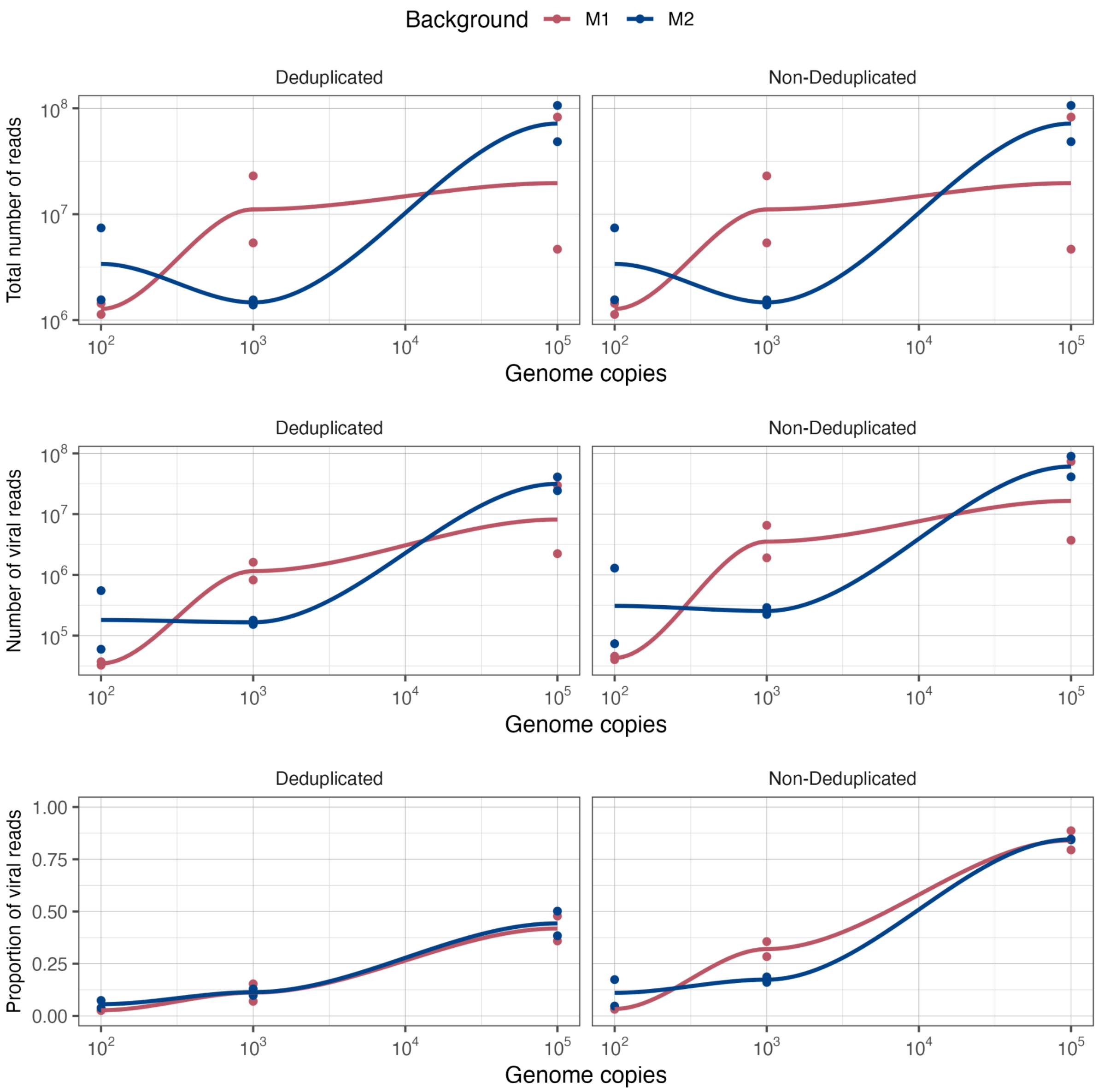

**Figure S4. Genome coverage of Human Adenovirus 40.** Number of reads (deduplicated) are shown for sites across the genome. The annotations on top of each plot reference the spike-in load and individual sample ID.

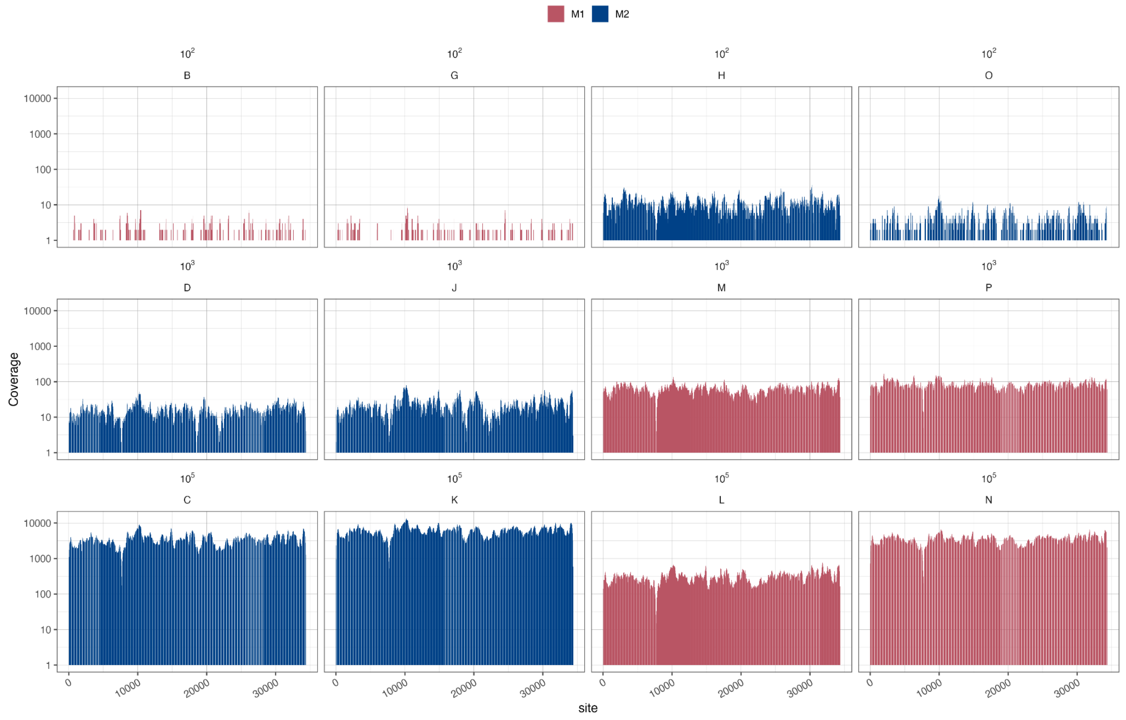

**Figure S5. Genome coverage of Human betaherpesvirus.** Number of reads (deduplicated) are shown for sites across the genome. The annotations on top of each plot reference the spike-in load and individual sample ID.

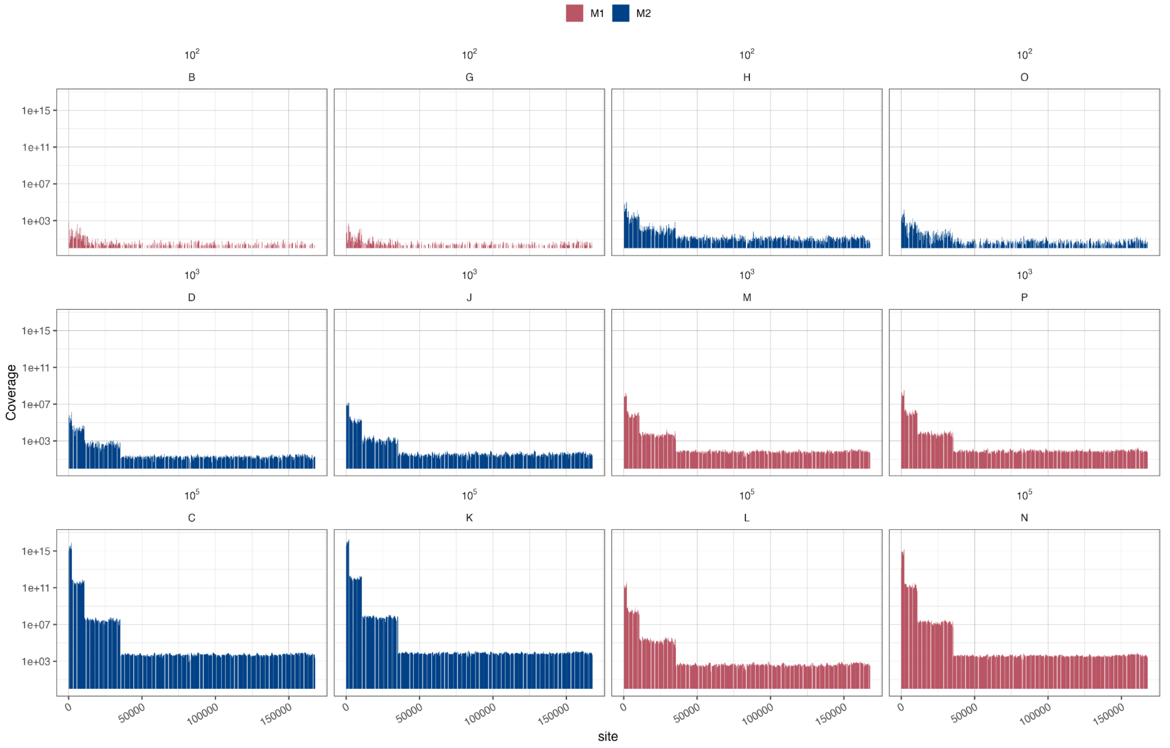

**Figure S6. Genome coverage of Human respiratory syncytial virus.** Number of reads (deduplicated) are shown for sites across the genome. The annotations on top of each plot reference the spike-in load and individual sample ID.

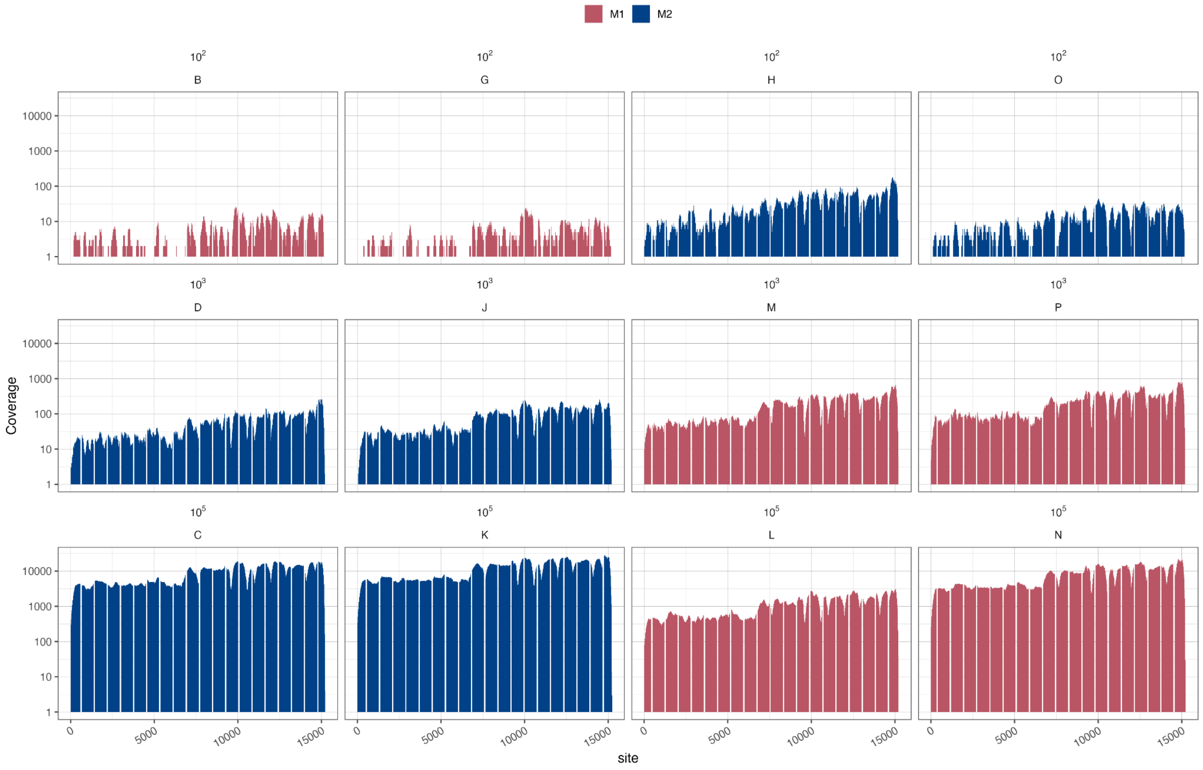

**Figure S7. Genome coverage of Influenza B virus.** Number of reads (deduplicated) are shown for sites across the genome. Segments are each row. The plots are now split by background and colors correspond to spike-in viral loads.

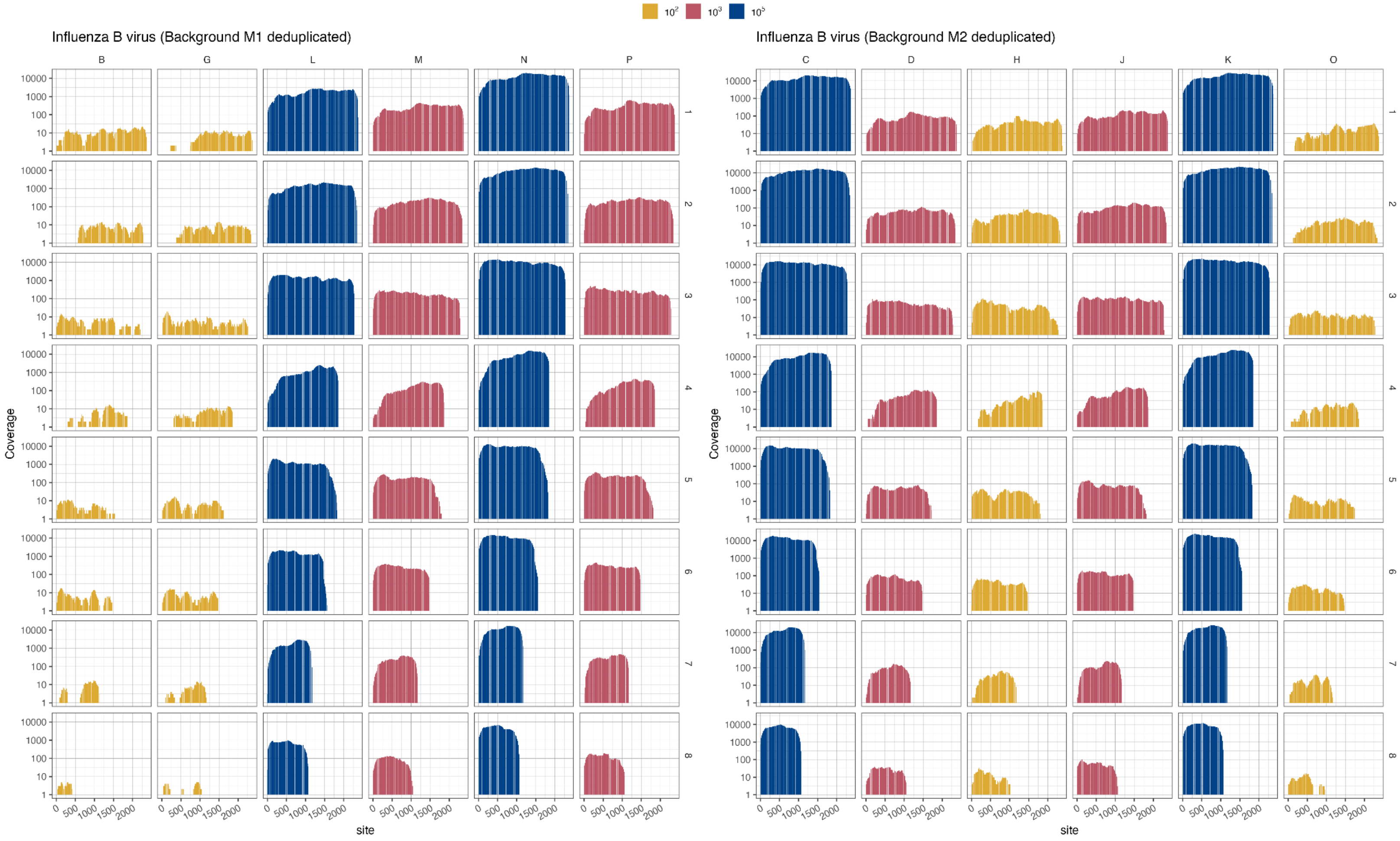

**Figure S8. Genome coverage of Mammalian orthoreovirus 3.** Number of reads (deduplicated) are shown for sites across the genome. Segments are each row. The plots are now split by background and colors correspond to spike-in viral loads.

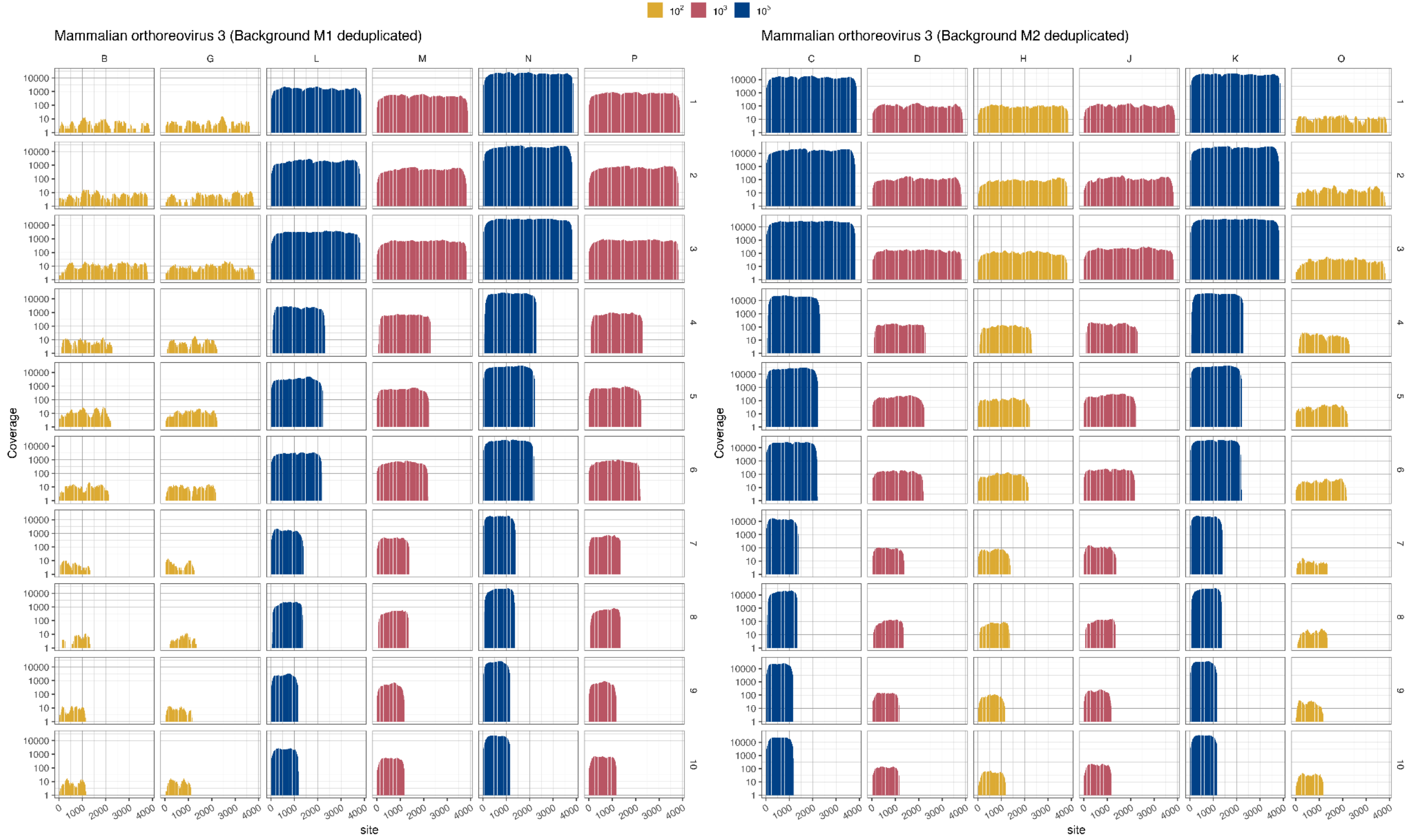

**Figure S9. Genome coverage of Zika virus.** Number of reads (deduplicated) are shown for sites across the genome. The annotations on top of each plot reference the spike-in load and individual sample ID.

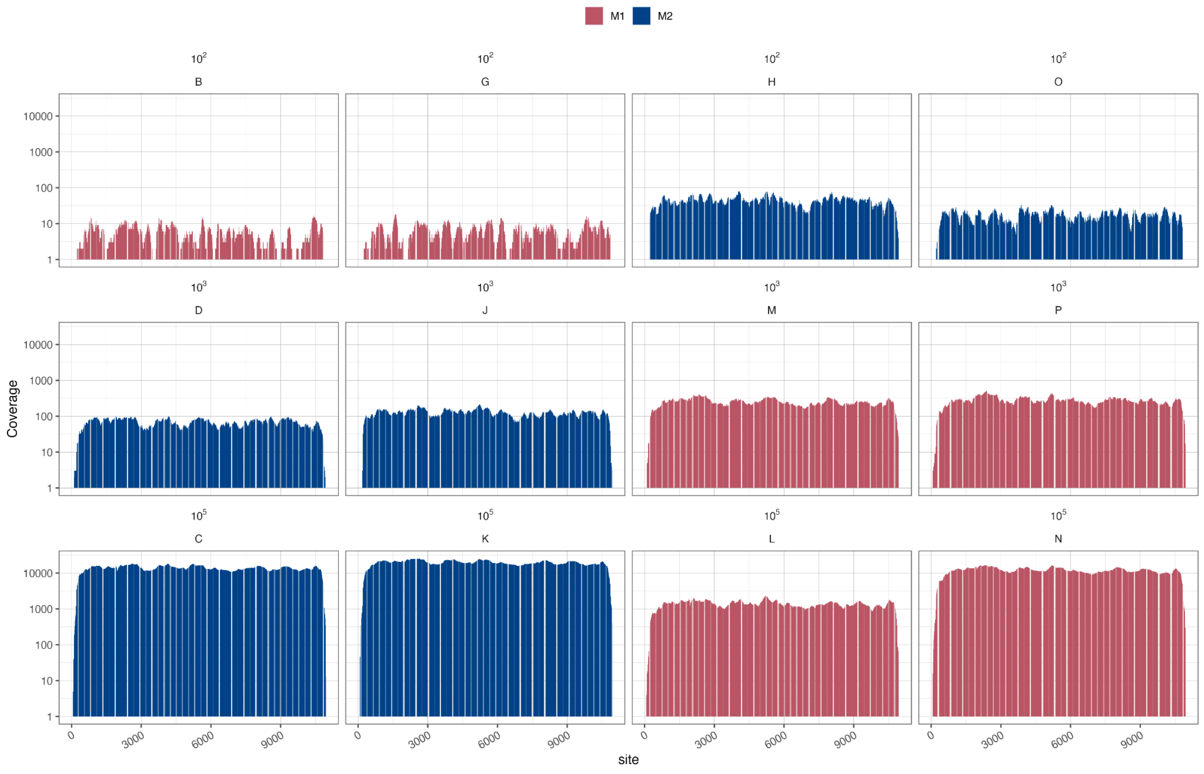

**Figure S10. Reanalyzed published data assessing read count and depth.** Reanalysis of Buddle and colleagues [1] data assessing spike-in loads of ATCC nucleic acid mix, which ranged from 60 to 6000 genome copies per ml.

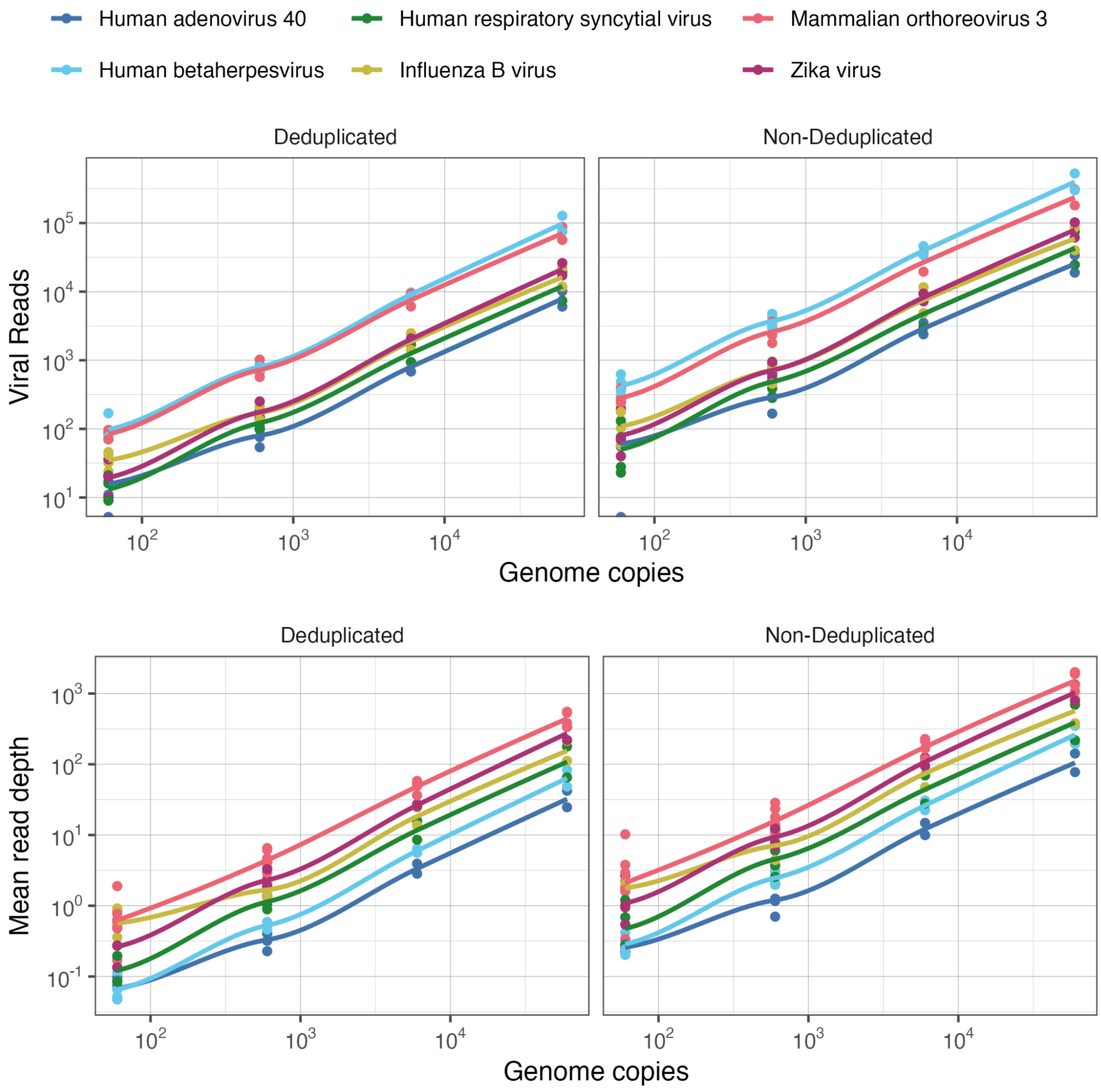

**Figure S11. Reanalyzed published data assessing genome coverage**. Reanalysis of Buddle and colleagues [1] data assessing spike-in loads of ATCC nucleic acid mix. The concentration of ATCC mix (genome copies/ml) are shown above each plot.

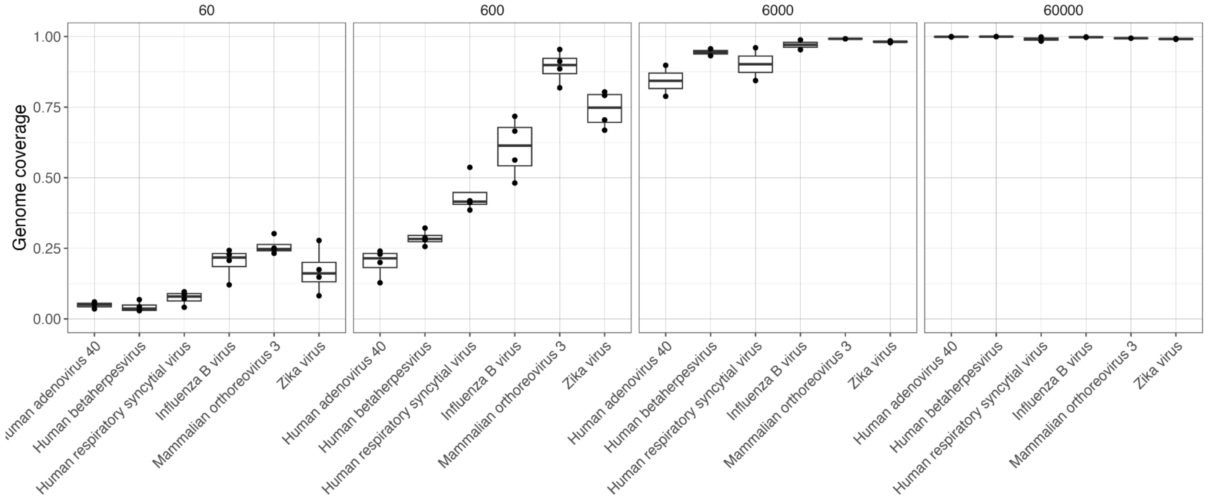

**Figure S12. Reanalyzed published data assessing Human Adenovirus 40.** Reanalysis of Buddle and colleagues [1] data assessing Adenovirus. Spike-in coverage (genome copies per ml) by rows - only two replicates were at higher spike-ins.

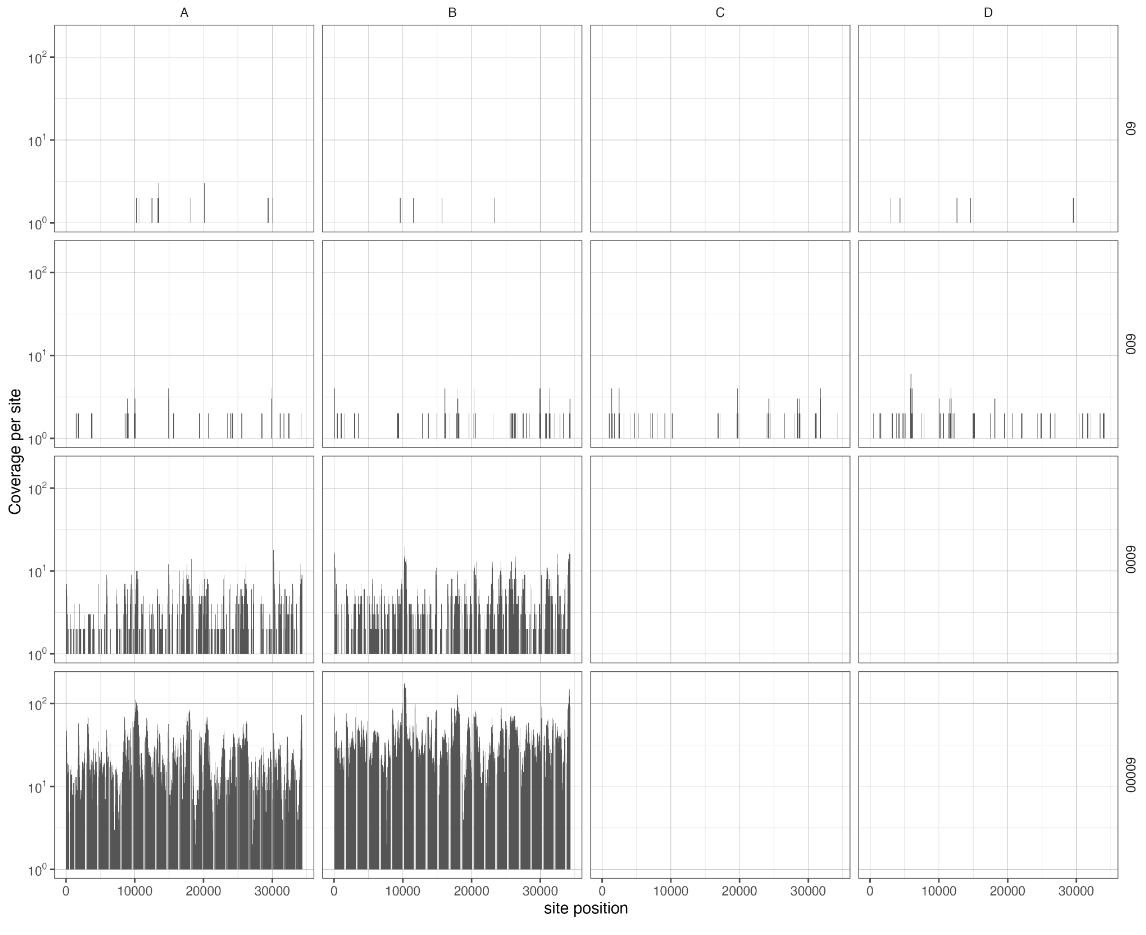

**Figure S13. *Kobuvirus* and *Cardiovirus* read depth.** Mean read depth is shown for both shotgun sequencing and target enrichment for both sample backgrounds (M1, M2). The spike-in virus concentrations are shown in different panels for target enriched sequencing data.

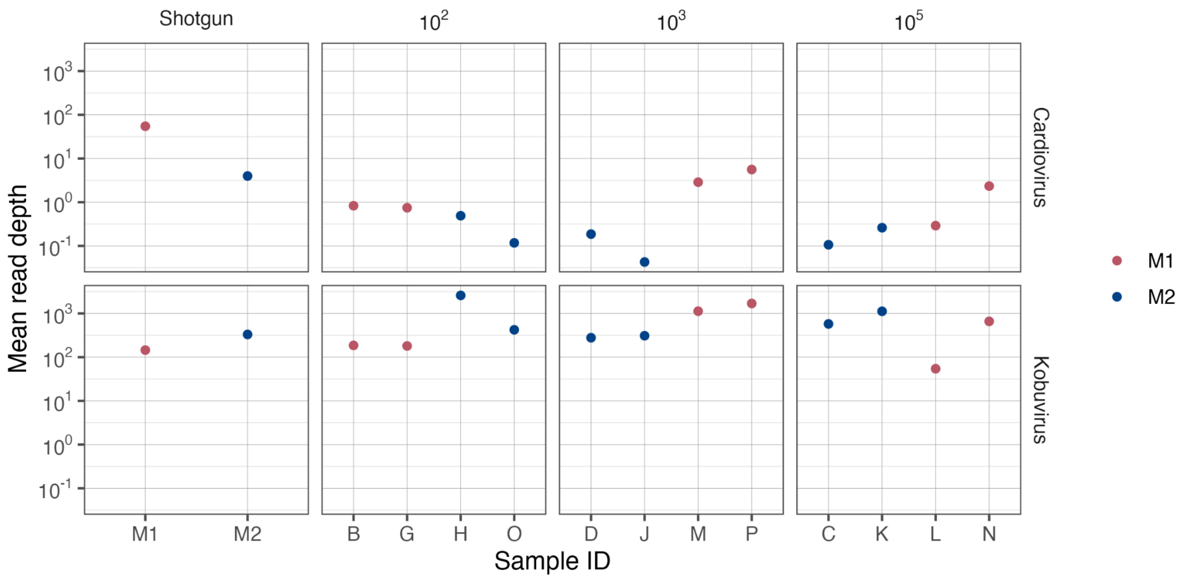

**FIgure S14. *Kobuvirus* and *Cardiovirus* genome coverage.** Proportion of genome covered for both shotgun sequencing and target enrichment for both sample backgrounds (M1, M2).The spike-in virus concentrations are shown in different panels for target enriched sequencing data.

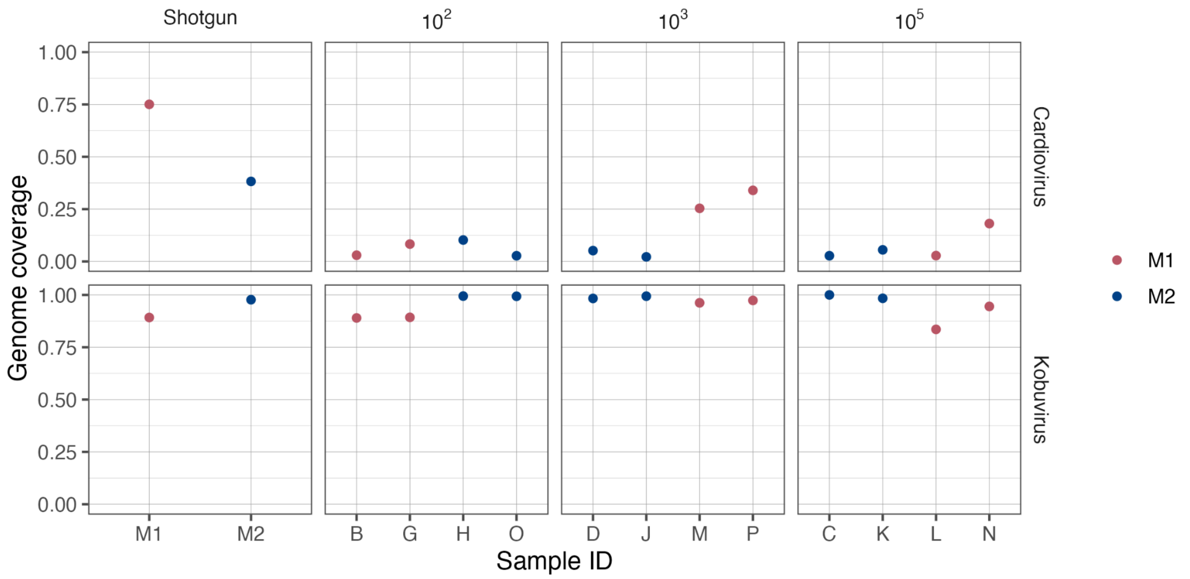

**Figure S15. *Cardiovirus* read depth across the genome.** Number of reads (deduplicated) are shown for sites across the genome. The annotations on top of each plot reference the spike-in load and individual sample ID or shotgun sequencing run.

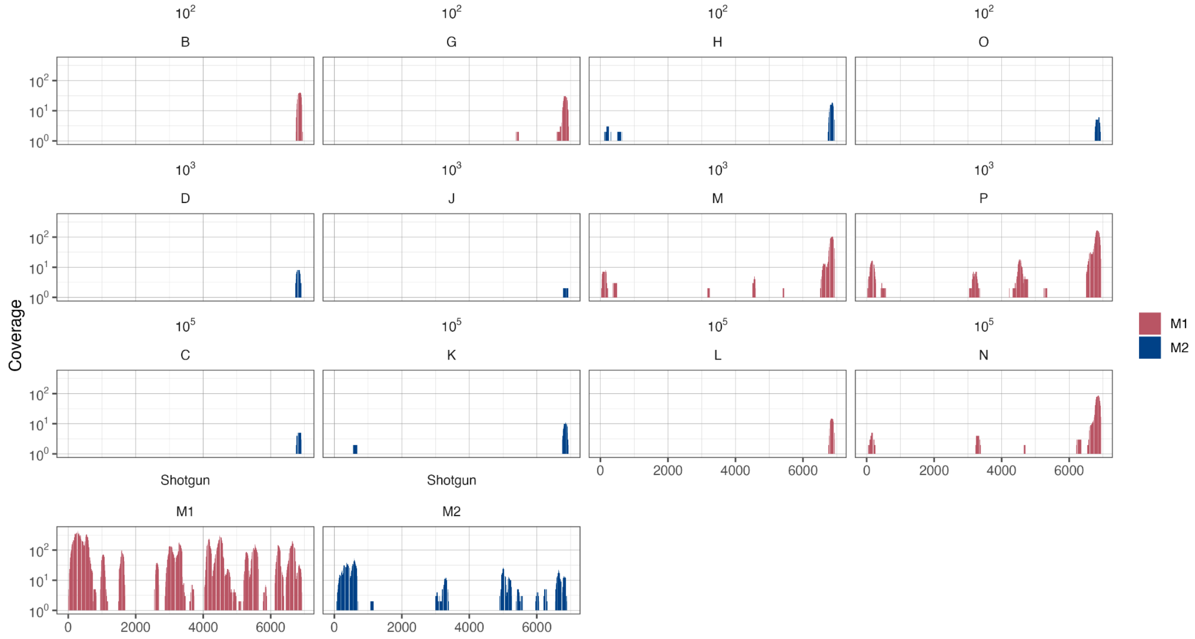

**Figure S16. *Kobuvirus* read depth across the genome.** Number of reads (deduplicated) are shown for sites across the genome. The annotations on top of each plot reference the spike-in load and individual sample ID or shotgun sequencing run.

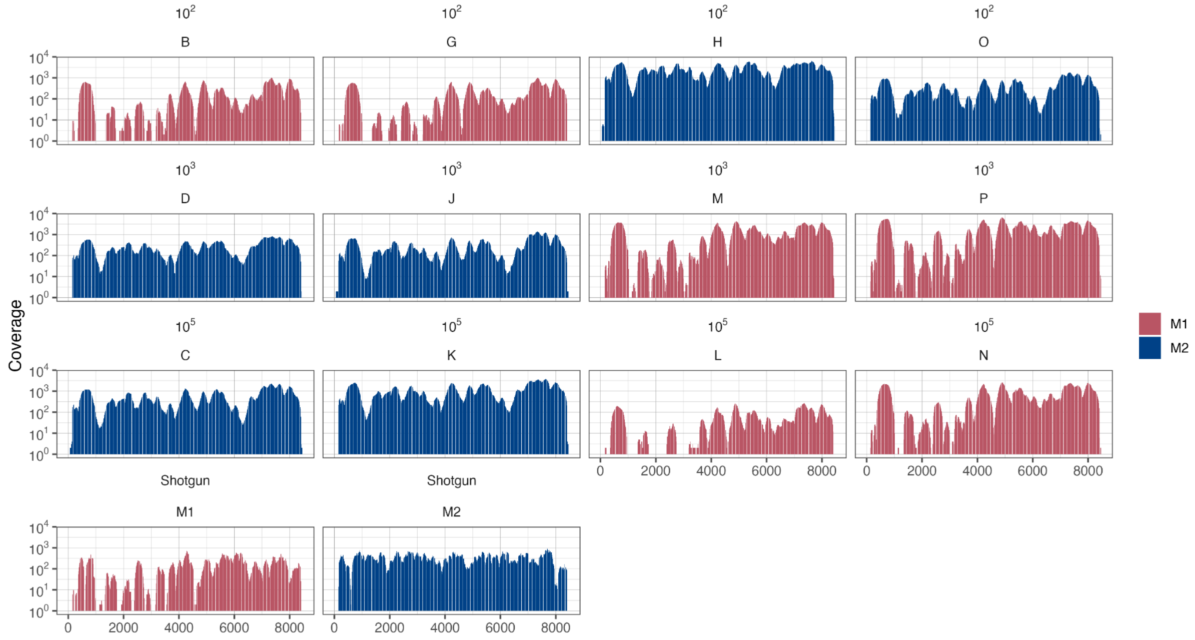
